# Phylogenomics places orphan protistan lineages in a novel eukaryotic super-group

**DOI:** 10.1101/227884

**Authors:** Matthew W. Brown, Aaron Heiss, Ryoma Kamikawa, Yuji Inagaki, Akinori Yabuki, Alexander K Tice, Takashi Shiratori, Ken-ichiro Ishida, Tetsuo Hashimoto, Alastair G.B. Simpson, Andrew J. Roger

## Abstract

Recent phylogenetic analyses position certain ‘orphan’ protist lineages deep in the tree of eukaryotic life, but their exact placements are poorly resolved. We conducted phylogenomic analyses that incorporate deeply sequenced transcriptomes from representatives of collodictyonids (diphylleids), rigifilids, *Mantamonas* and ancyromonads (planomonads). Analyses of 351 genes, using site-heterogeneous mixture models, strongly support a novel supergroup-level clade that includes collodictyonids, rigifilids and *Mantamonas*, which we name ‘CRuMs’. Further, they robustly place CRuMs as the closest branch to Amorphea (including animals and fungi). Ancyromonads are strongly inferred to be more distantly related to Amorphea than are CRuMs. They emerge either as sister to malawimonads, or as a separate deeper branch. CRuMs and ancyromonads represent two distinct major groups that branch deeply on the lineage that includes animals, near the most commonly inferred root of the eukaryote tree. This makes both groups crucial in examinations of the deepest-level history of extant eukaryotes.

## Introduction

Our understanding of the eukaryote tree of life has been revolutionized by genomic and transcriptomic investigations of diverse protists, which constitute the overwhelming majority of eukaryotic diversity (Burki 2014; Simpson and Eglt 2016). Phylogenetic analyses of supermatrices of proteins typically show a eukaryote tree consisting of five-to-eight ‘super-groups’ that fall within three even-higher-order assemblages: i) Amorphea (Amoebozoa plus Obazoa, the latter including animals and fungi), ii) Diaphoretickes (primarily Sar, Archaeplastida, Cryptista and Haptophyta), and iii) Excavata (Discoba and Metamonada) (Adl et al. 2012). Recent analyses (Derelle et al. 2015) place the root of the eukaryote tree somewhere between Amorphea and the other two listed lineages; Derelle *et al.* (2015) termed this the ‘Opimoda-Diphoda’ root. There is considerable debate over the position of the root, however (e.g., (Cavalier-Smith 2010; Katz et al. 2012; He et al. 2014))

Nonetheless, there remain several ‘orphan’ protist lineages that cannot be assigned to any super-group by cellular anatomy or ribosomal RNA phylogenies (e.g., (Brugerolle et al. 2002; Glücksman et al. 2011; Heiss et al. 2011; Cavalier-Smith 2013; Yabuki, Eikrem, et al. 2013; Yabuki, Ishida, et al. 2013)). Recent phylogenomic analyses including *Collodictyon*, *Mantamonas*, and ancyromonads indicate that these particular ‘orphans’ branch near the base of Amorphea (Zhao et al. 2012; Cavalier-Smith et al. 2014), the same general position as the purported Opimoda-Diphoda root. This implies, i) that these lineages are of special evolutionary importance, but also, ii) that uncertainty over their phylogenetic positions will profoundly impact our understanding of deep eukaryote history. Unfortunately their phylogenetic positions indeed remain unclear, with different phylogenomic analyses supporting incompatible topologies, and often showing low statistical support (Cavalier-Smith et al. 2014). This is likely due in part to the modest numbers of sampled genes for some/most species examined to date (Cavalier-Smith et al. 2014; Torruella et al. 2015). Therefore, we undertook phylogenomic analyses that incorporated deeply-sequenced transcriptome data from representatives of Collodictyonidae, *Mantamonas*, Ancyromonadida, and Rigifilida.

## Results

Using a custom phylogenomic pipeline plus manual curation we generated a dataset of 351 orthologs. The dataset was filtered of paralogs and potential cross-contamination by visualizing each protein’s phylogeny individually, then removing sequences whose positions conflicted with a conservative consensus phylogeny (as in (Tice et al. 2016; Kang et al. 2017)) (supplementary methods). We selected data-rich species to represent the phylogenetic diversity of eukaryotes. Our primary dataset retained 61 taxa, with metamonads represented by two short-branching taxa (*Trimastix* and *Paratrimastix*). We also analyzed a 64-taxon dataset containing three additional longer-branching metamonads. Maximum likelihood (ML) and Bayesian analyses were conducted using site-heterogeneous models; LG+C60+F+ Γ and the associated PMSF model (LG+C60+F+ Γ-PMSF) as implemented in IQ-Tree (Wang et al. 2017) and CAT-GTR+Γ in PhyloBayes-MPI, respectively. Such site-heterogeneous models are important for deep-level phylogenetic inference with numerous substitutions along branches (Lartillot et al. 2007; Le et al. 2008; Wang et al. 2008; Pisani et al. 2015; Wang et al. 2017).

Our analyses of both 61- and 64-taxon datasets robustly recover well-accepted major groups including Sar, Discoba, Metamonada, Obazoa, and Amoebozoa (Fig. 1, S1). Cryptista (e.g. cryptomonads and close relatives) branches with Haptophyta (Fig. 1) in the LG+C60+F+ Γ-PSMF analyses as well as in one set of two converged PHYLOBAYES-MPI chains under the CAT-GTR model (Fig. S2). However another pair of converged chains places Haptophyta as sister to Sar while Cryptista nests within Archaeplastida (Fig. S3), which is largely consistent with some other recent phylogenomic studies (e.g., (Burki et al. 2016)). Excavata was never monophyletic, with Discoba forming a clan with Diaphoretickes taxa (Sar, Haptophyta, Archaeplastida+Cryptista) and Metamonada grouping with Amorphea plus the four orphan lineages targeted in this study (see below). Malawimonads, which are morphologically similar to certain metamonads and discobids (Simpson 2003), also branch amongst the ‘orphans’ (see below).

**Figure 1.**
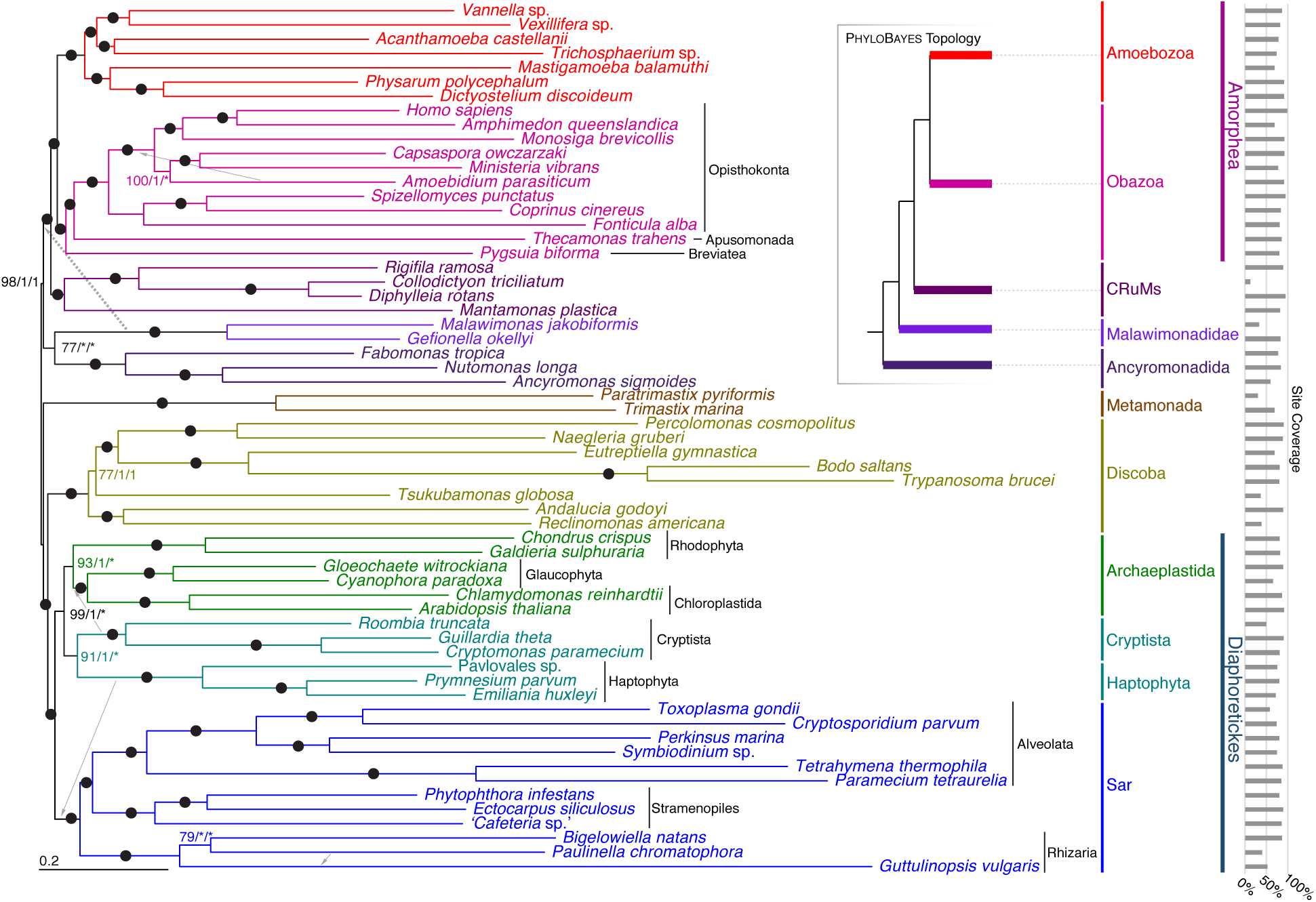
Phylogenetic tree for 61 eukaryotes, inferred from 351 proteins using Maximum Likelihood (LG+C60+F+ Γ-PMSF model). The numbers on branches show (in order) support values from 100 real bootstrap replicates (LG+C60+F+ Γ-PMSF model) and posterior probabilities both sets of converged chains in PHYLOBAYES-MPI under CAT-GTR+Γmodel (i.e., MLBS/PP/PP). Filled circles represent maximum support with all methods; asterisks indicate a clade not recovered in the PHYLOBAYES analysis. Further, the dashed arrow indicates the placement of malawimonads inferred with PHYLOBAYES-MPI (see also inset summary tree), and grey arrows indicate the placements of other lineages in the PHYLOBAYES-MPI analyses.

Phylogenies of both datasets place all four orphan taxa near the base of Amorphea (Fig. 1, Fig. S1). The uncertain position of the eukaryotic root (discussed above) therefore makes it unclear which bipartitions are truly clades, and which could be interrupted by the root. To allow efficient communication, we discuss the phylogenies as if the orphan taxa all lie on the Amorphea side of the root. We will also consider Amorphea as previously circumscribed (Adl et al. 2012): the least-inclusive clade or clan containing Amoebozoa and Obazoa.

Three of the orphan lineages are specifically related in our trees (Fig. 1, S1). In both 61-taxon and 64-taxon analyses, *Rigifila ramosa* (representing Rigifilida) forms a maximally-supported clade with the collodictyonids *Collodictyon triciliatum* and *Diphylleia rotans*. *Mantamonas plastica* then branches as their closest relative, with maximal support. This Collodictyonid+Rigifilida+*Mantamonas* clade (‘CRuMs’) forms the sister group to Amorphea, again with maximal support.

ML analyses and the converged PhyloBayes chains grouped ancyromonads, malawimonads, and CRuMs with Amorphea, with strong bootstrap support and Bayesian posterior probability (Fig. 1, 61 taxa; PMSF BS=98%, PP=1). Ancyromonads and malawimonads formed a clade in the ML analyses, but with equivocal support (Fig. 1, 61 taxa; BS=77%). Both sets of converged chains of the Bayesian analyses instead grouped malawimonads with CRuMs+Amorphea to the exclusion of ancyromonads (Fig. S2, S3, PP=1 for both); however some unconverged chains support an ancyromonad+malawimonad clade (data not shown). Lack of convergence amongst multiple chains using the CAT-GTR+Γ model is unfortunately common for large datasets, and often cannot be resolved by increasing the number of generations of Markov chain Monte Carlo within a reasonable time frame (Pisani et al. 2015; Kang et al. 2017). Instead we treat the two topologies recovered in these analyses as candidate hypotheses requiring further investigation.

We conducted approximately unbiased (AU) topology tests on the 61-taxon data set under the LG+C60+F+Γ mixture model (Table S1). These tests rejected the Phylobayes trees, as well as all trees optimized by enforcing constraints representing plausible alternative relative placements of ancyromonads, malawimonads, and metamonads.

The fastest evolving sites are expected to be the most prone to saturation and systematic error arising from model misspecification in phylogenomic analyses (Philippe et al. 2011). We conducted a ‘fast-site removal’ analysis with the 61-taxon data set and generated ultrafast bootstrap support (UFBOOT) values (Minh et al. 2013) for relevant groups as sites were progressively removed from fastest to slowest (Fig. 2A). All groups of interest receive reasonably strong support until ~44,000-48,000 sites were removed, when support fell markedly for the ancryomonad+*malawimonad* clade and the Amorphea+CRuMs+ancryomonad+Malawimonas clan. At this point, a notable proportion of the bootstrap trees show malawimonads and/or ancyromonads grouping with metamonads. This decline in support for the ancryomonad+malawimonad group reverses somewhat with further site removal, before support falls again as overall phylogenetic structure is lost when ~76,000 sites are removed (Fig. 2A).

**Figure 2.**
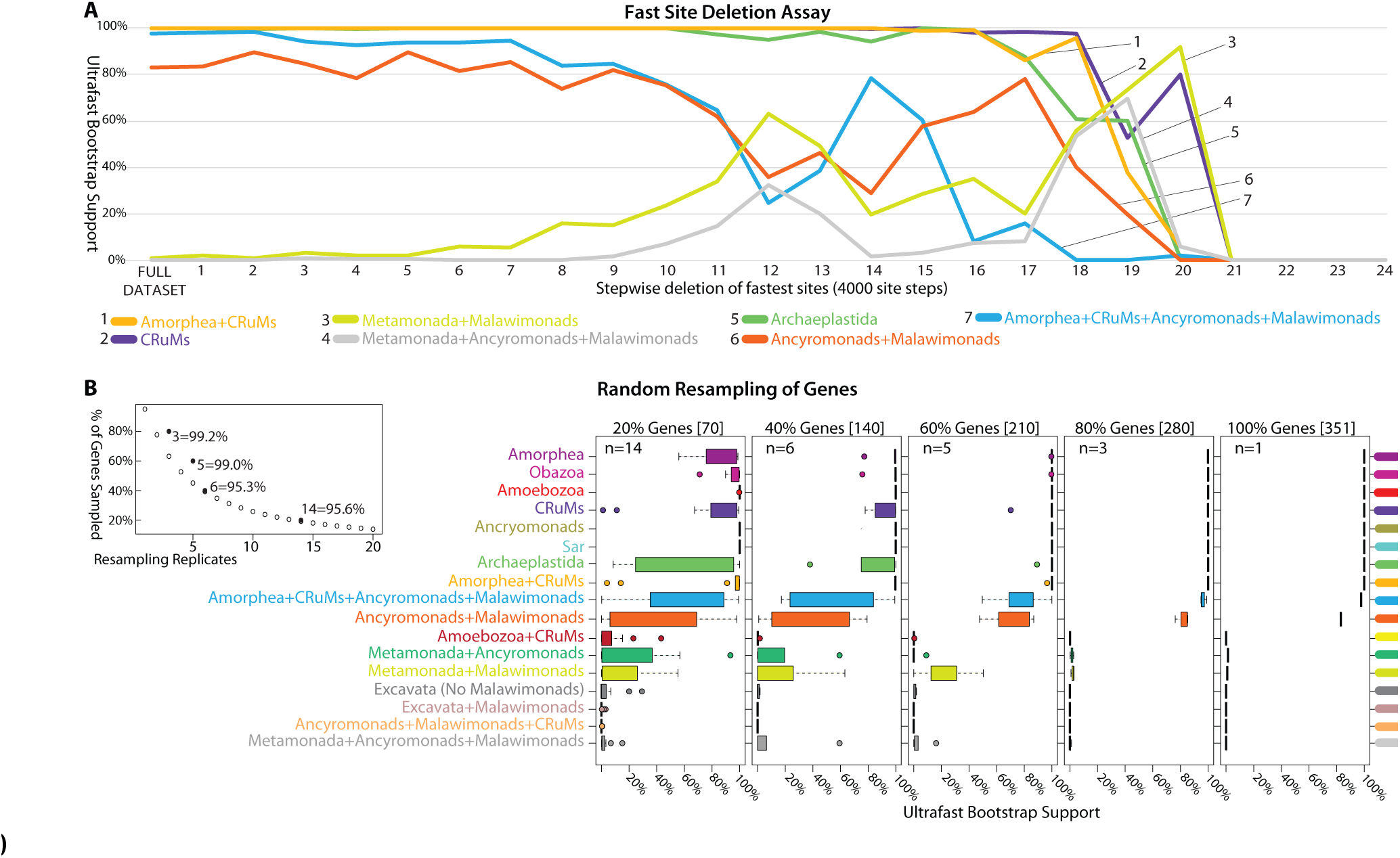
Effects of fast evolving sites and random subsampling of genes on our phylogenomic analyses. (A) Sites were sorted based on their rates of evolution estimated under the LG+F+Γ model and removed from the data set from highest to lowest rate. Each step has 4,000 of the fastest evolving sites removed progressively. The bootstrap values (UFBOOT; LG+C60+F+ Γ-PSMF model) for each bipartition of interest are plotted. (B, C) Effects of random subsampling of genes within the 351-gene dataset. The following bipartitions were examined but received nearly 100% support accross the fast site deletion assay (data not shown); Amorphea, Obazoa, Amoebozoa, Ancryomonads, and Sar. The following bipartitions were examined but received nearly 0% support across the fast site deletion assay (data not shown); Amoebozoa+CRuMs, Metamonada+Ancyromonads, Excavata (No Malawimonads), Excavata+Malawimonads, and Ancyromonads+Malawimonads+CRuMs. (B) Effects of random resampling of genes on the bipartitions of interest. Inset panel is the calculation of the number of replicates (*n*) necessary for a 95% probability of sampling every gene when subsampling 20, 40, 60 and 80% of genes using the formula: 0.95 = 1 – (1 – *x*/100) where *x* is the percentage of genes subsampled. UFBOOT support values for all nodes of interest with the variability of support values illustrated by box-and-whisker plots.

To evaluate heterogeneity in phylogenetic signals amongst genes (Inagaki et al. 2009), we also inferred phylogenies from subsamples of the 351 examined genes (61-taxon dataset; Fig. 2B,C). For each subsample 20-80% of the genes were randomly selected, without replacement, with replication as per Fig. 2B (giving a >95% probability that a particular gene would be sampled at each level), and UFBOOT support for major clades was inferred (Fig. 2C). The ‘80% retained’ replicates gave nearly identical results to the full dataset, indicating that there was little stochastic error associated with gene sampling at this level. Support for the CRuMs clade is almost always high when 40%+ of genes are retained, while subsamples containing 60% of genes still showed differing support for a ancyromonad-malawimonad clade (as opposed to, for example, malawimonads branching with metamonads).

## Discussion

Our 351 protein (97,002 AA site) super-matrix places several orphan lineages in two separate clades emerging between Amorphea and all other major eukaryote groups. All methods recover a strongly supported clade comprising the free-swimming collodictyonid flagellates, the idiosyncratic filose protist *Rigifila* (Rigifilida) and the gliding flagellate *Mantamonas*. This clade is resilient to exclusion both of fast-evolving sites and of randomly selected genes. It is also consistently placed as the immediate sister taxon to Amorphea. This represents the first robust estimate of the positions of these three taxonomically poor but phylogenetically deep clades. Previous phylogenomic analyses placed collodictyonids in various positions, such as sister to either malawimonads or Amoebozoa, but often with low statistical support (Zhao et al. 2012; Cavalier-Smith et al. 2014). Placements of *Mantamonas* have varied dramatically. A recent phylogenomic study recovered a weak *Mantamonas*+collodictyonid clade in some analyses, but other analyses in the same study instead recovered a weak *Mantamonas*+ancyromonad relationship (Cavalier-Smith et al. 2014), and SSU+LSU rRNA gene phylogenies strongly grouped *Mantamonas* with apusomonads (Glücksman et al. 2011; Yabuki, Ishida, et al. 2013). Our study decisively supports the first of these possibilities. This is the first phylogenomic analysis incorporating Rigifilida: Previous SSU+LSU rRNA gene analyses recovered a negligibly supported collodictyonid+rigifilid clade, but not a relationship with *Mantamonas* (Yabuki, Ishida, et al. 2013).

Overall, the hypotheses that i) collodictyonids, rigifilids and *Mantamonas* form a major eukaryote clade, and ii) this clade is sister to Amorphea, are novel, plausible, and evolutionarily important. No name exists for this putative super-group, and it is obviously premature to propose a formal taxon. We suggest the place-holding moniker ‘CRuMs’ (Collodictyonidae, Rigifilida, *Mantamonas*), which is euphonic and evokes the species-poor nature of these taxa.

Whether ancyromonads branch outside Amorphea or within it has been disputed (Paps et al. 2013; Cavalier-Smith et al. 2014). Our study strongly places ancyromonads outside Amorphea, more distantly related to it than are the CRuMs. Ancyromonads instead fall ‘amongst’ the excavate lineages (Discoba, Metamonada, and Malawimonadidae). Resolving the relationships amongst ‘excavates’ is extremely challenging (Hampl et al. 2009; Derelle et al. 2015), and this likely contributed to our difficulty in resolving the exact position of ancyromonads vis-à-vis malawimonads. A close relationship between ancyromonads and some/all excavates would be broadly consonant with the marked cytoskeletal similarity between *Ancyromonas* and ‘typical excavates’ (Heiss et al. 2011). Certainly, our study flags ancyromonads as highly relevant to resolving relationships amongst excavates.

Both candidate positions for ancyromonads place them at the centre of a crucial open question: locating the root of the eukaryote tree. As discussed above, the latest analyses (Derelle et al. 2015) locate the root between Discoba+Diaphoretickes (‘Diphoda’) and a clade including Amorphea, collodictyonids and malawimonads (‘Opimoda’). Our phylogenies show the ancyromonad lineage emerging close to this split. One of the two positions we recovered would actually place ancyromonads either as the deepest branch within ‘Diphoda’, or the deepest branch within ‘Opimoda’, or even as sister to all other extant eukaryotes. This demonstrates the profound importance of including ancyromonads in future rooted phylogenies of eukaryotes, using datasets optimized for this purpose.

## Materials and Methods

Details of experimental methods for culturing, nucleic acid extraction and Illumina sequencing are described in the supplemental text.

### Phylogenomic data set construction

A reference data set of 351 aligned proteins described in (Kang et al. 2017) was used as the starting point for the current analysis, from which 61 or 64 taxa representing diverse eukaryotes were selected (see Table S2). Extensive efforts were made to exclude contamination and paralogs, as described in the supplemental text. Poorly aligned sites were excluded using BMGE (Criscuolo and Gribaldo 2010), resulting in an alignment of 97,002 amino acid (AA) sites with less than 25% missing data for both 61- and 64-taxon datasets (Table S2).

### Phylogenomic tree inference

Maximum likelihood (ML) trees were inferred using IQ-Tree v. 1.5.5 (Nguyen et al. 2015). The best-fitting available model based on the Akaike Information Criterion (AIC) was the LG+C60+F+ Γ mixture model with class weights optimized from the dataset and four discrete gamma (Γ) categories. ML trees were estimated under this model for both 61- and 64-taxon data sets. We then used this model and best ML tree under the LG+C60+F+ Γ model to estimate the ‘posterior mean site frequencies’ (PMSF) model (Wang et al. 2017) for both 61 (Fig. 1) and 64 (Fig. S1) taxon data sets. This LG+C60+F+ Γ-PMSF model was used to re-estimate ML trees, and for a bootstrap analysis of the 61-taxon dataset, with 100 pseudoreplicates (Fig. 1). AU topology tests under the LG+C60+F+ Γ were conducted with IQ-Tree to evaluate whether trees recovered by the Bayesian analyses or alternative placements of the orphan taxa could be rejected statistically.

Bayesian inferences were performed using Phyliobayes-MPI v1.6j (Rodrigue and Lartillot 2014), under the CAT-GTR+Γ model, with four discrete Γ categories. For the 61-taxon analysis, 6 independent Markov chain Monte Carlo chains were run for ~4,000 generations, sampling every second generation. Two sets of two chains converged (at 800 and 2,000 generations, which were respectively used as the burnin), with the largest discrepancy in posterior probabilities (PPs) (maxdiff) < 0.05. The topologies of the converged chains are presented in Fig. S3 and S4 and are mapped upon Fig. 1. For the 64-taxon analysis, four chains were run for ~3,000 generations. Two chains converged at ~200 generations, which was used as the burnin, (maxdiff = 0) and the posterior probabilities are mapped upon the ML tree in Fig. S1.

### Fast-site removal and gene subsampling analyses

For fast site removal, rates of evolution at each site of the 61-taxon dataset were estimated with Dist_Est (Susko et al. 2003) under the LG model using discrete gamma probability estimation. A custom Python script was then used to remove fastest evolving sites in 4,000-site steps. Random subsampling of 20, 40, 60, or 80% of the genes in the 61-taxon dataset was conducted using a custom Python script, with the number of replicates as given in Fig 2B. In both cases each step or subsample was analyzed using 1,000 UFBOOT replicates in IQ-Tree under the LG+C60+F+r-PMSF model.

### Data availability

All new transcriptomic data have been deposited at the National Center for Biotechnology Information under BioProjects PRJNA401035, as detailed in Table S1. All single gene alignments, masked and unmasked, and phylogenomic matrices are available in supplemental file Brown_etal.2017.CRuMs.tgz

## Acknowledgements

The authors thank Tom Cavalier-Smith and Ed Glücksman (Oxford University) for supplying cultures strains B-70 (*Ancyromonas sigmoides*), NYK3C (*Fabomonas tropica*), and Bass1 (*Mantamonas plastica*). The part of this work conducted at Dalhousie University was supported by NSERC Discovery grants awarded to AGBS (298366-2014) and AJR (2016-06792) respectively. AJR also acknowledges the Canada Research Chairs program for support. This project was supported in part by the National Science Foundation (NSF) Division of Environmental Biology (DEB) grant 1456054 (athttp://www.nsf.gov), awarded to MWB. Mississippi State University’s High Performance Computing Collaboratory provided some computational resources. The part of this work conducted at the University of Tsukuba was supported by grants from the Japan Society for the Promotion of Science (JSPS; 15H05606 and 15K14591 awarded to RK, 23117006 and 16H04826 awarded to YI, 15H04411 awarded to KI, and 15H05231 to TH) and by the “Tree of Life” research project (Univ. Tsukuba).

